# Analytical performance and concordance with next-generation sequencing of a rapid multiplexed dPCR panel for the detection of actionable DNA and RNA biomarkers in non-small cell lung cancer

**DOI:** 10.1101/2023.05.04.539400

**Authors:** Kerri Cabrera, Jeffery Gole, Bryan Leatham, Lucien Jacky, Bradley A. Brown

## Abstract

**Background:** Over the last ten years, the discovery and FDA approval of targeted therapies for lung cancer has significantly improved patient survival rates. However, despite these improved survival rates, only 68% of patients receive molecular testing that results in assignment of targeted therapy ^1,2^. Barriers to timely access to biomarker information include no testing ordered^3^,high nucleic acid input requirements, and problematic turnaround time (TAT) by NGS (> 14 days)^4^.

Here we report the analytical performance and concordance with next-generation sequencing (NGS) of a highly-multiplexed research use only (RUO) panel using digital PCR (dPCR). The HDPCR NSCLC panel reports the status for variants (SNV, indels, and fusions) in eight actionable genes using amplitude modulation and multi-spectral encoding in dPCR^5^.

**Methods:** The panel’s analytical sensitivity and reactivity were determined using DNA and RNA extracted from formalin-fixed paraffin-embedded (FFPE) tissue spiked with plasmid DNA or in-vitro transcribed RNA. Concordance was established on 106 FFPE samples previously characterized using the Oncomine Precision Assay® or pathology results. Discordant resolution was resolved with Archer Fusionplex® and Variantplex® panels.

**Results:** The analytical sensitivity, reported as estimated mutant allele fraction (MAF), for DNA targets (*EGFR* exon 19 deletions, *EGFR* exon 20 insertions, *EGFR* S768I, *EGFR* L858R, *EGFR* T790M, *EGFR* L861Q, *BRAF* V600E, *EGFR* G719X, *ERBB2* exon 20 insertions and *KRAS* G12C) ranged from 0.8% – 4.9% with 40 ng of DNA input, and 2.4% to 10.9% with 15 ng of DNA input. For RNA fusion targets (*ALK, RET, ROS, NTRK* 1/2/3, and *MET* exon 14 skipping), the analytical sensitivity ranged from 24 - 150 copies with 5 ng of total RNA input. The population prevalence-based coverage ranged from 89.2% to 100.0% across targets and >99.0% in aggregate. The accuracy of the assay was >97% with respect to the comparator method.

## Introduction

An estimated 236,740 cases of lung and bronchus cancer were reported in the United States in 2022, with a five-year relative survival rate of only approximately 23%^6^. In the last five years, targeted therapies have increased three-year survival rates by 2.3-13.7%^5^. As of the publication of this article, with 32 FDA-approved targeted therapies, lung cancer is the solid tumor type with the most approved therapies targeted against driver mutations^3,7–9^. Despite these advances, recent studies indicate that for individuals with NSCLC, equity and accessibility are the main limitations for the use of targeted therapies.^1,4,7,10^. A study leveraging a database with commercial and Medicare claims from over 500,000 patients with NSCLC in the United States reported that approximately 50% of patients do not obtain full biomarker testing. Of the patients that did receive biomarker test results, 29% did not get the appropriate targeted therapy ^4,7^.The top reasons cited by oncologists for not testing all lung cancer patients include tissue or sample limitations, cost of and access to testing, long turnarounds for sequential gene testing, or having to send out for testing ^3,11,12,13^. One way to approach this problem would be to increase the timely availability of rapid biomarker testing that can be performed locally in the hospital setting with quick turnaround time.

Two common molecular biomarker testing modalities include next-generation sequencing (NGS) and sequential single-gene PCR. While NGS provides sequence information of entire genes and regions of the genome enabling comprehensive detection of multiple variants when present, there are also several critical challenges with sequencing-based approaches. One key challenge is time from collection to reported result; 13.1% of NGS testing had a turnaround time (TAT) greater than 14 days, which exceeds the TAT guideline established by the College of American Pathologists, IASLC, and the Association for Molecular Pathology^7^. In addition to lengthy TAT, approximately 22% of patients do not receive NGS results because of insufficient sample or poor sample quality^7,14^. NGS assays feature high complexity workflows and analysis that return many variants, some of which are not clinically actionable. In sharp contrast, single-gene PCR tests are manageable in terms of complexity, cost, actionable marker detection, and turnaround time. However, sample sufficiency remains a hurdle because of the need to split the sample across many tests or wells to get a complete result^15,16^.

In this article, we describe a highly multiplexed digital PCR (dPCR) assay that detects 15 relevant NSCLC variants in eight genes using amplitude modulation and multi-spectral encoding^17^. The panel is designed to capture actionable variants in NSCLC, is compatible with formalin-fixed paraffin embedded (FFPE) tissue specimens and utilizes a low mass input ^18–24^. The use of dPCR for the panel reduces the complexity of the workflow and TAT by reducing the number of user manipulations in comparison to NGS-based workflows. Cloud based analysis simplifies and accelerates results interpretation, allowing for results generation in less than 24 hours. Furthermore, testing for multiple actionable biomarkers in a single test potentially improves tissue requirements compared to single gene testing. The panel performance reported here includes analytical sensitivity, analytical reactivity (inclusivity), and method correlation with current NGS methodologies.

## Materials and Methods

### Materials

Plasmids containing the target sequences were obtained from IDT (San Diego, CA, USA). FFPE specimens were obtained from Precision for Medicine (Frederick, MD, USA), BioChain (Newark, CA, USA), or CHTN (Durham, NC, USA). Oncomine Precision Assay testing was performed by Precision for Medicine, and results were reported for all positive specimens. Negative specimens, non-tumor adjacent tissue, were reported negative by pathology. The mean age of the individual at time of sample acquisition was 63.5 years (standard deviation of 11.1).

### HDPCR NSCLC

High Definition (HDPCR) NSCLC panel utilizes dPCR, where endpoint fluorescent intensities are modulated such that each unique target produces a unique endpoint intensity ^17,25^. The HDPCR NSCLC Panel (ChromaCode, Carlsbad, CA) consists of three wells; two wells detect DNA targets, and one well detects RNA fusions. All runs were performed on the QIAcuity® (Qiagen, Germany) using the QIAcuity Nanoplate 26K 24-well plate. The master mix for DNA wells was formulated by combining 10.5 μL of QIAcuity Probe Master Mix, 8.4 μL of HDPCR Mix and 2.1 μL of molecular grade water per reaction. The master mix for each RNA well was formulated by combining 10.5 μL of QIAcuity OneStep Advance Probe Master Mix (Qiagen, Germany), 0.45 μL of OneStep RT Mix (Qiagen, Germany), 8.4 μL of HDPCR Mix, and 1.68 μL of molecular grade water per reaction. After preparation of the master mix, 21 μL of the sample was added to 21 μL of the appropriate master mix and mixed thoroughly. From this mixture, 39 μL was added to a well on the QIAcuity Nanoplate. The plate then underwent thermocycling on the QIAcuity according to the instructions for use. Analysis was carried out using ChromaCode Cloud, cloud-based analysis, which reported out detected targets and MAF. The MAF is calculated as (target counts / IC counts) x 100.

### Analytical Sensitivity

Negative FFPE background was prepared by extracting DNA and RNA from pathology-negative FFPE samples using the Maxwell HT FFPE DNA Isolation System. The limit of detection for DNA targets was established by spiking plasmids containing the target sequences into negative FFPE background at various MAF concentrations. The RNA fusion targets were transcribed from plasmids using the HiScribe T7 High Yield RNA Synthesis Kit (NEB, Ipswich, MA), isolated using the Monarch Kit (NEB, Ipswich, MA) and spiked into negative FFPE RNA background.

The limit of detection for DNA targets was established at two input amounts;20 ng and 7.5 ng of DNA per well (40 ng and 15 ng in total for both DNA wells). The RNA fusion targets were tested at 5 ng total RNA input. Range-finding was conducted by testing decreasing serial dilutions. For each target, the lowest concentration at which all replicates were positive during range-finding was evaluated with 20 replicates. The limit of detection is reported at the lowest concentration, where greater than 18/20 replicates were detected.

### Analytical Inclusivity

*In-silico* analysis of designs was performed using the COSMIC Mutation Database ^28^. The parameters utilized in filtering the data are recorded in **Supplemental 1**. After filtering, prevalence was calculated based on the count number of distinct entries in the “sample name field”. Analytical inclusivity was evaluated for the HDPCR NSCLC Panel by spiking in quantified plasmids containing the different sequences into the appropriate negative FFPE background (RNA or DNA, depending on the well). Each plasmid was tested at 3-5X the limit of detection in three replicates. If no replicate was detected, the plasmid was tested at 10X higher concentration. If any replicate was negative at the higher concentration, the assay was determined to be not inclusive for the specific sequence. Prevalence was estimated from the reported occurrences of unique Sample ids for each COSMIC ID (LEGACY_MUTATION_ID) associated with a reportable in the filtered COSMIC Mutation Database described in **Supplemental 1**.

### Concordance Study

106 unique FFPE blocks (77 positive samples with the Oncomine Precision Assay and 29 pathology negative samples) from lung tissue were enrolled in the study. DNA and RNA were extracted from a single 10 μm curl using the Maxwell HT FFPE DNA Isolation System (Promega, Madison, WI) on the KingFisher™ Flex instrument (Thermofisher, Carlsbad, CA). Following extraction, eluates were quantified using the Qubit dsDNA BR Assay Kit (Invitrogen, Waltham MA) or the Qubit RNA BR Assay Kit (Invitrogen, Waltham, MA). Samples were evaluated with the HDPCR NSCLC Panel according to the above mentioned methods. Results from the Oncomine Precision Assay (from separate sections of the same block) and the HDPCR NSCLC Panel was compared, and any discordant samples (same section as evaluated by HDPCR) were sent for discordant resolution using the Variant Plex solid tumor focus (Archer, Boulder, CO) or the Fusion Plex Lung (Archer, Boulder, CO). Results from discordant analysis were then detailed for each target.

## 3 Results

### Analytical Sensitivity, Limit of Detection (LOD)

The analytical sensitivity is reported in estimated Mutant Allele Fraction (MAF). Each assay features an internal control (IC) to determine if sufficient amplifiable nucleic acid has been loaded into the well. At 20 ng input (1854 average IC Counts) DNA per well, the LOD ranged from 0.8% to 4.9% MAF (**Table 1**). When the total DNA input was decreased to 7.5 ng per well (476 average IC Counts), the limit of detection ranged from 2.4% to 10.9% MAF (**Table 1**). With an input of 5 ng (97 average IC counts), the limit of detection for RNA targets ranged from 24 to 150 counts (**Table 2**). These results indicate that even with minimal inputs of DNA and RNA, the HDPCR NSCLC panel is sensitive for all targets.

**Table 1:** Limit of Detection for DNA targets. The limit of detection reported in estimated MAF as determined by ChromaCode Cloud software. Input amount is defined in ng for each well as measured by Qubit. Mean counts represent the average number of positive partitions at the reported limit of detection. The detection rate is the positive results out of 20 total replicates.

|  | 7.5 ng Input |  |  | 20 ng Input |  |  |
| --- | --- | --- | --- | --- | --- | --- |
| Target | Mean Counts | Detection Rate | MAF | Mean Counts | Detection Rate | MAF |
| *EGFR* exon 19 deletions | 76.7 | 20/20 | 10.9% | 30.2 | 20/20 | 1.2% |
| *EGFR* G719X | 40.9 | 20/20 | 6.2% | 33.9 | 20/20 | 2.7% |
| *EGFR* S768I | 51.0 | 20/20 | 9.6% | 67.2 | 20/20 | 3.2% |
| *ERBB2* insertions | 41.6 | 20/20 | 7.3% | 32.2 | 20/20 | 2.4% |
| *EGFR* L858R | 71.5 | 20/20 | 8.8% | 56.0 | 19/20 | 2.6% |
| *EGFR* exon 20 insertions | 60.4 | 20/20 | 10.1% | 58.3 | 20/20 | 4.9% |
| *BRAF* V600E | 21.0 | 20/20 | 2.4% | 45.1 | 20/20 | 2.2% |
| *KRAS* G12C | 22.5 | 20/20 | 3.6% | 34.2 | 20/20 | 2.3% |
| *EGFR* T790M | 37.3 | 20/20 | 9.5% | 19.5 | 20/20 | 0.8% |
| *EGFR* L861Q | 40.5 | 20/20 | 8.3% | 40.7 | 20/20 | 3.5% |

**Table 2:**
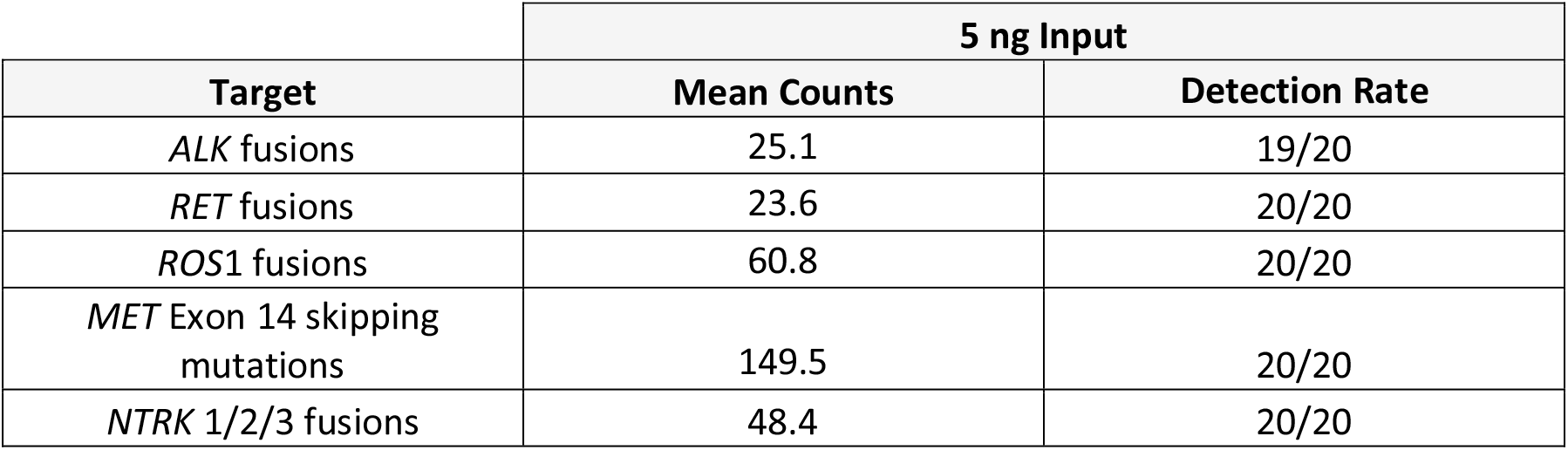
Limit of Detection for RNA Targets. The limit of detection reported mean counts by ChromaCode Cloud software. Input amount is defined in ng for each well as measured by Qubit. Mean counts represent the average number of positive partitions at the reported limit of detection. The detection rate is the positive results out of 20 total replicates.

### Inclusivity

Inclusivity was evaluated both *in silico* for DNA targets, and empirically, for DNA and RNA fusion targets, by testing plasmids spiked in FFPE negative matrix. The *in-silico* analysis resulted in 31 DNA targets flagged for empirical evaluation. The results for *in silico* and empirical analysis for DNA targets are reported in **Table 3**. RNA fusion targets were all evaluated empirically, with 96 different variations evaluated. The results are reported in **Table 4** and **Supplemental 2**.

**Table 3:** Inclusivity Results for DNA targets. In silico and empirical (bench testing) results for inclusivity by target. “Evaluated” represents the number of sequences (unique COSMIC IDs) evaluated in the study. “Not Inclusive” are the sequences that are not detected by the panel. “Detected” are the sequences that are detected by the panel.

|  | In Silico Analysis |  |  | Empirical Analysis |  | In Silico + Empirical | Prevalence per 1000 |  |
| --- | --- | --- | --- | --- | --- | --- | --- | --- |
| Target | Evaluated | Not Inclusive | Inclusive | Evaluated | Detected | Prevalence Coverage | Positive Sample per 1000 | Missed Calls due to Inclusivity |
| *EGFR* Exon 20 Insertions | 24 | 5 | 19 | 8 | 8 | 89.2% | 9 | 1 |
| *EGFR* Exon 19 Deletions | 23 | 2 | 13 | 16 | 14 | 98.6% | 131 | 1.8 |
| *EGFR* G719X | 3 | 3 | 3 | 3 | 3 | 100% | 10 | 0 |
| *ERBB2* | 6 | 1 | 5 | 5 | 5 | 99.4% | 9 | <0.1 |

**Table 4:** Inclusivity Results for RNA Targets. Empirical (bench testing) results for inclusivity by target. “Evaluated” represents the number of sequences (unique COSMIC IDs) evaluated in the study. “Not Inclusive” are the sequences that are not detected by the panel. “Detected” are the sequences that are detected by the panel.

| Target | Evaluated | Detected | Overall Inclusivity<br>(Prevalence Weighted) | Positive<br>Patients per<br>1000 | Missed Calls<br>due to<br>Inclusivity |
| --- | --- | --- | --- | --- | --- |
| *ALK* | 41 | 40 | 99% | 27 | <0.3 |
| *ROS1* | 25 | 25 | 100% | 4 | 0 |
| *RET* | 20 | 20 | 98.7% | 4 | <0.1 |
| *NTRK* | 10 | 10 | 95.7% | 0 | 0 |

### Concordance Study

The HDPCR NSCLC panel was used to evaluate 106 unique FFPE samples. The IC in DNA wells failed in 15 of the 106 samples, while the RNA IC failed in 6 of the 106 samples. The failure of the IC in samples was correlated with the source vendor, which ranged from 0% to 63% internal control failures (**Supplemental 3**).

In samples where the IC passed, the concordance of the HDPCR NSCLC panel with the comparator method was 97.8% (1399/1430). The positive percent agreement (PPA) for individual targets ranged from 50.0%-100.0% and the positive predictive value (PPV) ranged from 62.5 to 100.0% before discordant resolution, low values are driven by the low number of positives samples available for some targets and high level of discordant results. The negative percent agreement (NPA) ranged from 97.0%-100.0% and the negative predicted value (NPV) ranged from 94.3%-100% (**Table 5**). Discordant samples were evaluated, from the same extraction, if possible, with either the VariantPlex solid tumor focus panel for DNA targets or the FusionPlex Lung for RNA targets. After discordant resolution, each target’s PPA ranged from 71.4%-100.0% and PPV ranged from 71.4%-100.0%. The NPA ranged from 97.9%-100% and the NPV ranged from 97.9%-100% for individual targets after discordant resolution.

**Table 5:** Concordance data reported by target for Oncomine Precision Assay and HDPCR NSCLC Panel. True Positive (TP), True Negative (TN), False Positive (FP), Positive Percent Agreement (PPA), Positive Predictive Value (PPV), Negative Predictive Value (NPV) and Negative Percent Agreement (NPA). Adjusted values after discordant resolution are placed in parenthesis. ^1^ Two *BRAF* false negatives were not detected by discordant resolution. One *BRAF* false positive was detected with discordant resolution.. ^2^ One *EGFR* Exon 19 Deletion false negative was not detected in discordant resolution, the QC score was below threshold for discordant resolution. One *EGFR* Exon 19 Deletion false positive was detected by discordant resolution. ^3^ Two *EGFR* L858R false negative were not detected by discordant resolution. Three *EGFR* L858R false positives were detected in discordant resolution. ^4^ Two *EGFR* T790M false negative were not detected by discordant resolution. One *EGFR* T790M false positive was detected by discordant resolution. ^5^ One *EGFR* L861Q false positive was detected by discordant resolution. ^6^ One ERBB false positive was detected with discordant resolution. ^7^ One *KRAS* G12C false positive was detected by discordant resolution. ^8^ Two of the three *ALK* false negatives were not detected by discordant resolution. One of the two detected a novel *NTRK* fusion, but failed to pass the QC threshold. One of the three false negative was detected by discordant resolution and reported as a novel fusion. ^9^ Two of four *NTRK* false negative were not detected by discordant resolution. Additionally, two of the four *NTRK* false negative were detected by discordant resolution but was reported as a novel fusion. ^10^ Three of three *RET* false negatives were not detected by discordant resolution. ^11^ Two of the three *ROS*1 false positives were detected by discordant resolution. One *ROS*1 false positive was detected by discordant resolution and reported as a novel variant.

| Target | TP | TN | FP | FN | PPA (PPA post discordant) | NPA (NPA post discordant) | PPV (PPV post discordant) | NPV (NPV post discordant) |
| --- | --- | --- | --- | --- | --- | --- | --- | --- |
| *BRAF* V600E<sup>1</sup> | 8 | 84 | 1 | 2 | 80.0%<br>(90.0%) | 98.8%<br>(100.0%) | 88.9%<br>(100.0%) | 97.7%<br>(98.8%) |
| *EGFR* exon 19 deletion<sup>2</sup> | 3 | 90 | 1 | 1 | 75.0%<br>(100.0%) | 98.9%<br>(100.0%) | 75.0%<br>(100.0%) | 98.9%<br>(100.0%) |
| *EGFR* L858R<sup>3</sup> | 5 | 85 | 3 | 2 | 71.4%<br>(100.0%) | 96.6%<br>(100.0%) | 62.5%<br>(100.0%) | 97.7%<br>(100.0%) |
| *EGFR* S768I | 3 | 92 | 0 | 0 | 100.0%<br>(100.0%) | 100.0%<br>(100.0%) | 100.0%<br>(100.0%) | 100%<br>(100.0%) |
| *EGFR* T790M<sup>4</sup> | 3 | 89 | 1 | 2 | 60.0%<br>(100.0%) | 98.9%<br>(100.0%) | 75.0%<br>(100.0%) | 97.8%<br>(100.0%) |
| *EGFR* G719X | 2 | 90 | 0 | 0 | 100.0%<br>(100.0%) | 100.0%<br>(100.0%) | 100.0%<br>(100.0%) | 100%<br>(100.0%) |
| *EGFR* exon 20 insertion | 7 | 84 | 1 | 0 | 100.0%<br>(100.0%) | 98.8%<br>(98.8%) | 87.5%<br>(87.5%) | 100%<br>(100.0%) |
| *EGFR* L861Q<sup>5</sup> | 0 | 91 | 1 | 0 | N/A<br>(100%) | 98.9%<br>(100.0%) | N/A<br>(100.0%) | 100%<br>(100.0%) |
| *ERBB2* insertions<sup>6</sup> | 8 | 83 | 1 | 0 | 100.0%<br>(100.0%) | 98.8%<br>(100.0%) | 88.9%<br>(100.0%) | 100%<br>(100.0%) |
| *KRAS* G12C<sup>7</sup> | 3 | 88 | 1 | 0 | 100.0%<br>(100.0%) | 98.9%<br>(100.0%) | 75.0%<br>(100.0%) | 100%<br>(100.0%) |
| *ALK* fusions<sup>8</sup> | 5 | 91 | 0 | 3 | 62.5%<br>(71.4%) | 100.0%<br>(100.0%) | 100.0%<br>(56.6%-100.0%) | 96.8%<br>(91.05-98.9%) |
| *MET* exon14 skipping | 4 | 94 | 1 | 0 | 100%<br>(100.0%) | 98.9%<br>(98.9%) | 80.0%<br>(80.0%) | 100%<br>(100%) |
| *NTRK1/2/3* fusions<sup>9</sup> | 0 | 95 | 0 | 4 | N/A<br>(N/A) | 100.0%<br>(100.0%) | N/A<br>(N/A) | 96.0%<br>(98.0%) |
| *RET* fusions<sup>10</sup> | 3 | 93 | 0 | 3 | 50.0%<br>(100.0%) | 100.0%<br>(100.0%) | 100%<br>(100.0%) | 96.9<br>(100.0%) |
| *ROS1* fusions<sup>11</sup> | 4 | 92 | 3 | 0 | 100%<br>(100.0%) | 96.8%<br>(97.9%) | 57.1%<br>(71.4%) | 100%<br>(100.0%) |

In total, there were 31 discordant results with the comparator method. Discordant resolution agreed with the HDPCR NSCLC Panel in 71.0% (22/31) of discordant results and aligned with the comparator in 29.0% (9/31) of results. Of the nine discordant results that aligned with the comparator method, 44.4% (4/9) of the results were novel fusions that are outside the inclusivity of the HDPCR NSCLC Panel, with 5/9 incongruous samples remaining. All discordant analysis results are described in **Table 6**.

**Table 6:**
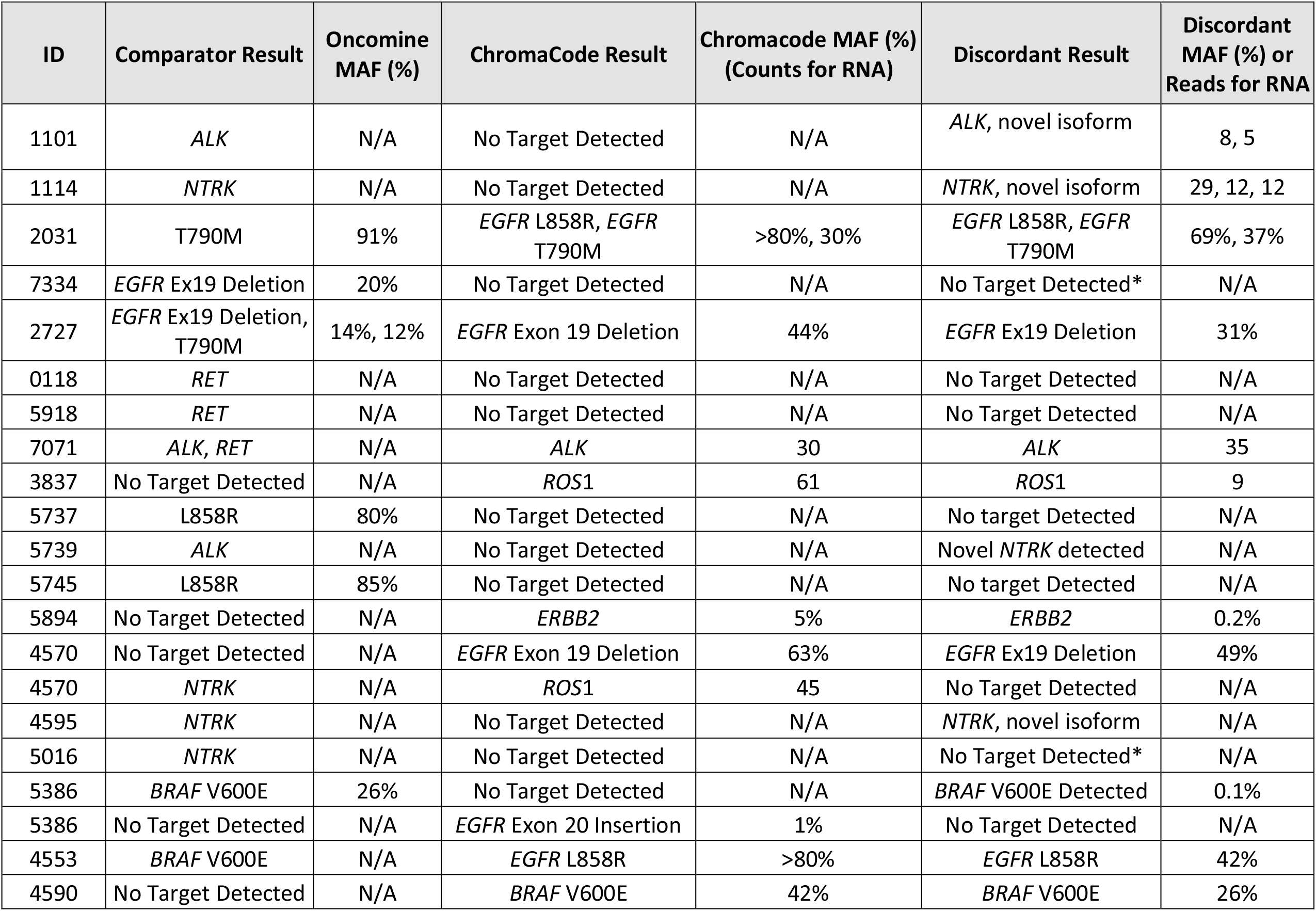
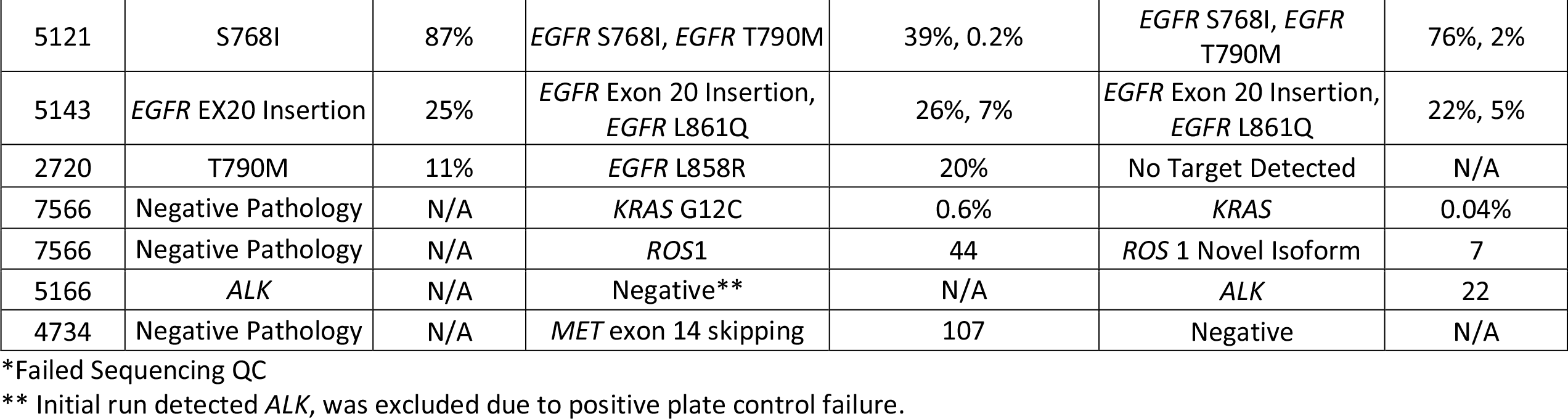
Results from discordant analysis. Results for all discordant samples with comparator and discordant test results. MAF is reported for all samples where a DNA target was detected. RNA counts, number of positive partitions, are reported for all RNA detections with the ChromaCode panel. Fusion partners for discordant resolution are reported in **Supplement 4**.

## 5 Conclusions and Discussions

The HDPCR NSCLC Panel uses a more simplified workflow involving only 2 touch points post extraction on dPCR, allowing a TAT of less than 4 hours excluding extraction time. The results from the studies presented here illustrate that the HDPCR NSCLC Panel utilizing dPCR achieved sensitivity down to 0.8% MAF and greater than 99% accuracy with comparator results after discordant resolution.

Lung samples are often characterized by limited tissue availability for molecular testing. Collection methods for specimens in NSCLC include fine needle aspirates (FNA), core needle biopsy (CNB), and resected tissue, offering varying amounts of surveyable genetic material. The input amounts can lead to 6.4-22% “no call” or quantity not sufficient (QNS) rates which increase as ng input decreases for NGS-based tests^14,26^. dPCR has been demonstrated to have sensitivity with low DNA input amounts but is limited by the scope of the variants examined in a single well^15,16^. Here we demonstrate a MAF limit of detection between 2.4-10.9% at as little as 15 ng of DNA input split across two wells (7.5 ng per well). While at high DNA input amounts, we report a MAF limit of detection between 0.8-4.9% at 40 ng input (20 ng per well). In the future, the HDPCR NSCLC panel can be evaluated with cfDNA and cfRNA to potentially provide a more sensitive and straightforward workflow for liquid biopsy applications.

One disadvantage of traditional PCR approaches to the detection of variants is poor sequence coverage or inclusivity^27^. The HDPCR NSCLC Panel was designed for high inclusivity for highly variable targets like *EGFR* exon 20 insertions (89%), *EGFR* exon 19 deletions (95%), and RNA fusions (95-100%). The high inclusivity provides increased confidence in negative results.

The HDPCR NSCLC Panel demonstrated high concordance (>97%) with the comparator methods. Discordant results, 71.0% (22/31), between the comparator and the HDPCR NSCLC panel resolved in favor of HDPCR. Taken together, the results demonstrate how the HDPCR NSCLC Panel can test for actionable variants with low nucleic acid input using a simple PCR-based workflow. For the RNA fusion targets, the primary source of false negatives, five of nine, was due to the detection of novel fusions that are not within the scope of the HDPCR panel.

The ability to provide faster, accurate results for actionable biomarkers is a key step toward democratizing testing. Here we present a dPCR panel that provides a coverage of actionable biomarkers with high concordance with NGS. The simplified workflow and analysis make it a potential solution for improving accessibility to relevant biomarker testing.

